# Varying water deficit stress (WDS) tolerance in grain amaranths involves multifactorial shifts in WDS-related responses

**DOI:** 10.1101/153577

**Authors:** América Tzitziki González-Rodríguez, Ismael Cisneros Hernández, Norma A. Martínez-Gallardo, Erika Mellado-Mojica, Mercedes López-Pérez, Enrique Ramírez-Chavez, Jorge Molina-Torres, John P. Délano-Frier

## Abstract

In this study, water deficit stress (WDS)-tolerance in several cultivars of grain amaranth species (*Amaranthus hypochondriacus* [Ahypo], *A. cruentus* [Acru] and A. *caudatus* [Acau]), in addition to *A. hybridus* (Ahyb), an ancestral amaranth, was examined. Ahypo was the most WDS-tolerant species, whereas Acau and Ahyb were WDS-sensitive. Data revealed that the differential WDS tolerance observed was multifactorial. It involved increased proline and raffinose (Raf) in leaves and/ or roots. Higher foliar Raf coincided with induced *Galactinol synthase 1* (*AhGolS1*) and *Raffinose synthase* (*AhRafS*) expression. Unknown compounds, possibly larger RFOs, also accumulated in leaves of WDS-tolerant amaranths, which had high Raf/ Verbascose ratios. Distinct nonstructural carbohydrate (NSC) accumulation patterns were observed in tolerant species under WDS and recovery, such as: i) high Hex/ Suc ratios in roots coupled to increased cell wall and vacuolar invertase and sucrose synthase activities; ii) a severer depletion of starch reserves; iii) lower NSC content in leaves, and iv) higher basal hexose levels in roots which further increased under WDS. WDS-marker gene expression patterns proposed a link between amaranth’s WDS tolerance and abscisic acid-dependent signaling. Results obtained also suggest that *AhTRE*, *AhTPS9*, *AhTPS11*, *AhGolS1 and AhRafS* are reliable gene markers of WDS tolerance in amaranth.

**Highlight:** Differential water deficit stress tolerance in grain amaranths and their ancestor, *Amaranthus hybridus*, is a multifactorial process involving various biochemical changes and modified expression patterns of key stress-related genes.

## Introduction

Plants have evolved to avoid, escape or tolerate stress conditions using numerous mechanisms that include several morphological, physiological and metabolic adaptations (Golldack *et al.*, 2014). Plant drought and salt adaptation involves control of water flux and cellular osmotic adjustment via the regulation of stomatal aperture, biosynthesis of osmoprotectants and reestablishment of the cellular redox status via the removal of reactive oxygen species (ROS) (Golldack *et al.*, 2014). Gene expression is also profoundly modified upon salt and drought stress. Stress-related genes code for proteins involved in osmolyte biosynthesis, detoxifying processes and transport, as well as in regulatory processes. Transcription factors (TFs), protein kinases, and phosphatases are central players in the latter. Both abscisic acid (ABA)-dependent and ABA-independent signaling pathways are activated to cope with abiotic stress (Krasensky and Jonak, 2012; Golldack *et al.*, 2014).

The accumulation of compatible solutes constitutes a protective mechanism employed by plants to ameliorate the damaging effects of drought and other abiotic stresses. This diverse group includes proline (Pro), soluble non-structural carbohydrates (NSCs; i.e., sucrose, glucose and fructose), and raffinose family oligosaccharides (RFOs), among others. They can accumulate in many plants in response to different stresses, and may perform roles other than osmoprotection, such as ROS scavenging or protein stabilization. Pro accumulation has also been found to promote plant recovery from drought stress (An *et al.*, 2013). Starch can be rapidly mobilized to provide soluble sugars. Thus, starch catabolism is accelerated in response to salt drought and other stresses usually as an osmotic adjustment response via increased soluble NSCs accumulation (Castrillón-Arbeláez *et al.*, 2012; Vargas *et al.*, 2013; Reguera *et al.*, 2013). These can maintain cell turgor or protect membranes and proteins from stress-related damage. Sucrose metabolism, via invertases, also regulates abiotic stresses responses by providing hexoses, as essential metabolites and signaling molecules (Ruan *et al.*, 2012) or promoting heat shock protein accumulation (Liu *et al.*, 2013). Likewise, glucose metabolism may prevent cell death via augmented reducing power and concomitant antioxidants biosynthesis (Bolouri-Moghaddam *et al.*, 2010). On the other hand, RFOs are extensively distributed in higher plants, functioning in carbon (C) storage and redistribution. (Ayre *et al.*, 2013). They are also known to accumulate during seed desiccation and/ or in leaves of plants subjected to various abiotic stresses, although their precise role in plant stress tolerance acquisition is not fully understood (Nishizawa *et al.*, 2008; ElSayed *et al.*, 2014). Biosynthesis of RFO originates from galactinol (Gol) generated from myo-inositol (MI) and UDP-galactose by Gol synthase (GolS). Gol subsequently acts as a galactose unit donor to Suc to generate raffinose (Raf), stachyose (Sta) and higher order RFOs, via their respective glycosyltransferases (Peterbauer and Richter, 2001).

Trehalose (Tre) is a non-reducing disaccharide present in trace amounts in most plants. It is presumably involved in the regulation of plant development and abiotic stress resistance. Thus, targeted manipulation of trehalose-6-phosphate (T6P), Tre’s precursor, also accumulating in trace amounts in most plants, has been found to improve abiotic stress tolerance and yield in some crop plants (Figueroa and Lunn, 2016). This phosphorylated precursor is synthesized by trehalose-6-phosphate synthases (TPSs) and may be subsequently dephosphorylated to Tre by trehalose-6-phosphate phosphatases (TPPs). Tre itself may be hydrolyzed to two glucose moieties by trehalase (TRE) (Lunn *et al.*, 2014). T6P’s role as a sensor of C availability has been proposed to involve a negative interaction with sucrose non-fermenting related kinase-1 (SnRK1) a known inhibitor of plant growth (Liu *et al.*, 2013; Delorge *et al.*, 2014; Lunn *et al.*, 2014; Tsai and Gazzarrini, 2014; Figueroa and Lunn, 2016).

The genus Amaranthus consist of 60-70 species. Some are consumed as vegetables or are used as a source of grain. The latter (*Amaranthus hypochondriacus*, *A. cruentus*, and *A. caudatus*) possess desirable agronomic characteristics and produce highly nutritional seeds. Moreover, they adapt easily to drought and poor soils (Caselato-Sousa and Amaya-Farfán, 2012). Domesticated grain amaranths presumably descend from wild *A. hybridus*, although their origin and taxonomic relationships are still uncertain (Sogbohossou and Achigan-Dako, 2014).

The physiological traits that enable amaranths to thrive in harsh conditions, such as drought, and be amenable for cultivation on marginal lands unsuitable for cereals, have been partly uncovered. In this work, we compared four amaranth species that differed in their tolerance to water-deficit stress (WDS) in order to identify common and/ or divergent responses to this condition. The changes in the expression, in leaves and roots, of RFO-biosynthetic genes and of genes involved in Tre metabolism and signaling were also evaluated. The content of RFOs, NSCs and Pro as well as invertases, sucrose synthase and amylase activity was also determined. The combined results of this study demonstrated that the differential WDS tolerance detected in the amaranth species tested, was the result of a multifactorial response

## Materials and Methods

### 2.1 Plant material

Three semi-domesticated grain amaranth species (*A. hypochondriacus* [Ahypo], *A. cruentus* [Acru] and *A. caudatus* [Acau]) (Sauer, 1967) together with an undomesticated vegetable amaranth (*A. hybridus* [Ahyb]), believed to be grain amaranths’ ancestor (Stetter and Schmid, 2017), were employed in the greenhouse experiments here described. All plant materials were provided by Dr. Eduardo Espitia Rangel, INIFAP, México, curator of the Mexican amaranth germplasm collection. Approximately 3 week-old plants having 9-10 expanded leaves were employed for experimentation. These were grown in 1.3 L plastic pots containing 250 g of a general substrate in a conditioned growth chamber, as described previously (Délano-Frier *et al.*, 2011). A total of 8 cultivars/ accessions of at least one of the above species was tested, as follows: Ahypo (“Gabriela”, “Revancha” and “DGTA” cultivars); Acru (“Amaranteca”, Dorada” and “Tarasca” cultivars), Acau (no classification available) and Ahyb (accession N°. 1330).

### 2.2 Water-deficit (WDS) stress experiments

All WDS experiments were performed in a commercial green house with zenithal and lateral type ventilation (Baticenital 850; ACEA S.A., Mexico) in May to August of 2015. The average temperatures in the greenhouse ranged between 15°C (night) and 38°C (day), with an average 55% R.H. The experiments were performed under natural light and photoperiod (≈ 1300 μE, ≥ 12 h light). An initial experiment was performed to screen the 8 cultivars/ accessions mentioned above for their tolerance to WDS. WDS was established by withholding watering for 7 or 10 days, time after which moderate to severe plant wilting was evident. WDS tolerance was scored by determining the leaf water potentials at the end of the WDS treatments and the percentage of recovery one day after normal watering was restored following stress (results not shown). This led to the selection of the 4 materials for subsequent experimentation which were the following: Ahypo (var. “Gabriela”) and Acru (var. “Amaranteca”), classified as “WDS tolerant”, and Acau and Ahyb, as “WDS susceptible”.

Subsequently, two tandem experiments were performed in the above conditions to test WDS tolerance based on soil water depletion. Prior to the start of the WDS trials, each experimental 1.3 L pot was weighed individually until maximum soil water retention capacity (SWC) was attained. WDS trials were started when all pots were at 90% SWC. Control plants were kept in these conditions for the duration of the experiments, whereas WDS was established by withholding watering until the SWC in each pot reached either 30% SWC (“moderate WDS, [MWDS]”) or 10% SWC (“severe WDS, [SWDS]”). These stress levels were reached approximately 5-6 and 9-10 days after regular watering was withheld, respectively. An additional group of plants was re-watered after reaching 10% SWC and was allowed to recover for 24 h (“recovery”, [R]). Weighing of the pots to determine water loss was done on a daily basis, taking care to ensure it was consistently done at approximately the same time of the day. Twelve plants having 10-to-12 expanded leaves were used per treatment. Once the desired conditions were reached, roots and leaves from 3 similarly treated plants were sampled and combined. Control plants were similarly sampled, generating four subsamples per experimental group. All pooled tissue samples were flash frozen in liquid N2 and stored at -70°C until needed.

### 2.3 Extraction of total RNA and gene expression analysis by RT-qPCR

Quantitative gene expression analysis using SYBR Green detection chemistry (Bio-Rad, Hercules, CA, USA) was performed as described previously (Palmeros-Suárez *et al.*, 2015). Primers design for the amplification of the pertinent amaranth gene transcripts employed a published methodology (Thornton and Basu, 2011) and was based on recently published genomic data (Clouse *et al.*, 2016) (Table S1). Relative gene expression was calculated using the comparative cycle threshold method (Livak and Schmittgen, 2001) using the *AhACT7*, *AhEF1α* and *AhβTub5* genes for data normalization.

### 2.4 Determination of NSC and Pro

Leaf and root samples collected from control plants, and from plants subjected to MWDS, SWDS or R, were used to quantify soluble NSCs and Pro contents, according to Palmeros-Suárez *et al.* (2015).

### 2.5 Determination of RFOs by HPAEC–PAD

Identification and determination of RFOs content in leaf and root samples was performed by High-Performance Anion-Exchange Chromatography with Pulsed Amperometric Detection (HPAEC–PAD), according to Mellado-Mojica *et al.* (2016). All chemicals used for the optimization of the chromatographic separation conditions and for quantitation were acquired from Sigma (Sigma-Aldrich, St. Louis, MO, USA),). These were the following: MI, Gol, Raf, Sta, and verbascose (Ver).

### 2.6 Determination of trehalose by GC/MS and thin layer chromatography (TLC) analysis

Tre levels were determined by GC/ MS using a ion selective method as described previously (Orona-Tamayo *et al.*, 2013). RFO analysis by TLC was performed using HPTLC silica gel 60 F254 plates as described previously (Waksmundzka-Hajnos *et al.*, 2008).

### 2.7 Determination of invertases, sucrose synthase and amylase activities

Vacuolar, cell wall, and cytoplasmic invertases and sucrose synthase (SuSy) activities were determined as described in Wright *et al.* (1998). Amylase activity was determined according to Bernfeld (1955). All assays were modified to fit a micro-plate format.

### 2.8 Statistical analysis

All experiments were conducted using a randomized complete block design. One-way ANOVAs were utilized to evaluate differences between treatment means. For ANOVAs where the F test was significant at *P* ≤ 0.05, the Tukey-Kramer test was applied. Statistical analysis was performed with R software (Development Core Team, https://www.r-project.org/).

## 3. Results

The initial screening to determine possible differences in WDS tolerance between amaranth species revealed that Ahypo cv. Gabriela, followed by Acru cv. Amaranteca were the most WDS tolerant species (with ca. 60% and 45% recovery after WDS, respectively). On the other hand, Acau was the most susceptible (with a 35% recovery rate to MWDS but unable to tolerate SWDS). Interestingly, completely desiccated Ahyb plants recovered from SWDS after watering was restored (results not shown).

A gene expression analysis in roots and leaves of the four amaranth genotypes was performed next. These were originally detected in a previous grain amaranth transcriptomic analysis (Délano *et al.*, 2011), and were later found to respond to severe defoliation in grain amaranth (Cisneros, 2016). Several genes involved in Tre biosynthesis and breakdown, including one class I TPS gene (*AhTPS1*), three TPP genes (*AhTPPA*, *AhTPPD* and *AhTPPI*) and one TRE gene (*AhTRE*) were analyzed. Also included were several non-catalytic class II TPS genes (*AhTPS5*, *AhTPS7*, *AhTPS8*, *AhTPS9*, *AhTPS10* and *AhTPS1 1*). Only the class I *TPS* gene (*AhTPS-AHYPO 004431*) and four additional *AhTPP* genes, annotated in the *A. hypochondriacus* genome (Clouse *et al.*, 2016), were not included in this study, whereas all 6 class II *TPS* genes were incorporated. All genes were named according to the closest homology shown with their respective *Arabidopsis thaliana* orthologs (Supplementary Fig. S1-S3). Also included were four genes involved in RFOs biosynthesis (Gol synthase [*AhGolS1* and *AhGolS2*], Raf synthase [*AhRafS*], and Sta synthase [*AhStaS*]) and a number of sucrose non-fermenting related kinases similarly shown to be affected by severe defoliation in grain amaranth (*SnRAK*, *SnRK1a*, *SnRK2.1* and *SnRK2.2*) (Cisneros, 2016). Finally, the expression of four ABA-related stress marker genes (*AhRAB18*, *AhABI5*, *AhDREB2C* and *AhLEA14*) were included as controls.

WDS had almost no effect on the expression of the class I *AhTPS1* gene in leaves (Table 1A). Only limited induction was detected in Ahyb and Acau. Similarly, the expression of class II *AhTPS5*, *AhTPS7* and *AhTPS8* in leaves of all plants was predominantly unaffected by stress or downregulated. Downregulation of these genes by MWDS in Ahypo was prominent. In contrast, *AhTPS11* responded strongly to WDS and R in practically all plants tested. This response was particularly evident during SWDS. The expression of the other two class II TPS genes was also induced chiefly during SWDS, although differences between species were observed in other conditions (Table 1B). Likewise, the *TPP* and *TRE* genes were, in general, unaffected or repressed by WDS in leaves. Noticeable exceptions were the induction, by WDS, of *AhTPPD* and *AhTPPI*, and the repression, in R, of *AhTPPA* and *AhTPPD*, in Ahypo. WDS also induced *AhTPPA, AhTPPI* and *AhTPPD* in Acru and Acau (Table 1C). Finally, *AhTRE* was exclusively induced by WDS in leaves of Ahypo (Table 1D).

**Table 1.** Relative expression values^1^ of genes involved in trehalose synthesis and degradation in leaves of four *Amaranthus* species subjected to two levels of water-deficit stress (moderate [M] and severe [S]) and to subsequent recovery ([R]). Induced (normalized expression values ≥ 2.0; in normal text) and repressed (normalized expression values ≤ 0.5; in italicized text) expression values are emphasized in bold.

| Gene | *A. hypochondriacus* |  |  | *A. cruentus* |  |  | *A. caudatus* |  |  | *A. hybridus* |  |  |
| --- | --- | --- | --- | --- | --- | --- | --- | --- | --- | --- | --- | --- |
|  | M | S | R | M | S | R | M | S | R | M | S | R |
| *AhTPS1* | 1.348 | 1.088 | 0.886 | 0.764 | 0.788 | 0.899 | 1.100 | 1.205 | **2.532** | 1.315 | **1.704** | **3.030** |
| *AhTPS5* | *0.233* | *0.268* | *0.272* | 0.552 | 0.526 | *0.343* | 0.664 | 0.606 | 0.645 | *0.423* | 0.536 | 0.638 |
| *AhTPS7* | **0.498** | 0.768 | 0.748 | 0.879 | 1.208 | 0.806 | 0.673 | 1.172 | **2.245** | 0.659 | 1.076 | 1.199 |
| *AhTPS8* | *0.250* | 0.902 | *0.235* | 0.967 | 0.896 | 0.711 | 1.076 | **1.835** | **1.621** | *0.386* | 0.827 | 0.974 |
| *AhTPS9* | 0.796 | **2.917** | *0.287* | 0.893 | **2.454** | 0.781 | **2.045** | **2.715** | **2.832** | 1.115 | **2.136** | 1.021 |
| *AhTPS10* | *0.291* | 1.016 | *0.264* | **1.890** | **2.589** | **6.992** | **2.382** | 1.262 | **2.892** | *0.438* | **1.651** | 1.154 |
| *AhTPS11* | **5.006** | **17.995** | 1.449 | **2.052** | **3.522** | **1.530** | **3.650** | **17.420** | **2.927** | **5.047** | **23.795** | **2.449** |
| *AhTPPA* | 0.515 | 1.121 | **1.971** | 1.189 | 0.856 | 0.534 | 1.170 | **1.514** | 1.054 | 0.751 | 1.491 | 1.083 |
| *AhTPPD* | *0.334* | *0.186* | **2.972** | **1.579** | 0.824 | 0.730 | **1.556** | 0.876 | 0.580 | 0.532 | 1.318 | **1.948** |
| *AhTPPI* | *0.297* | *0.279* | 0.769 | 1.022 | 0.591 | *0.369* | 0.951 | **1.573** | 1.062 | 0.628 | 1.176 | 1.016 |
| *AhTRE* | **2.047** | **2.170** | 1.276 | 0.583 | *0.324* | *0.474* | 0.562 | 0.643 | 1.143 | 0.996 | 0.716 | 0.987 |

<sup>1</sup>Calculated according to the comparative cycle threshold method (Livak and Schmittgen, 2001) using the *AhACT7*, *AhEF1a* and *AhβTub5* amaranth genes for data normalization.

The expression pattern of these genes changed in roots. The frequency with which they were induced in response to WDS or R was lower and their expression levels were reduced compared to those in leaves (Table 2). Thus, class I *AhTPS1* was repressed in WDS-tolerant species and unaltered in the other two (Table 2A). Likewise to leaves, class II *AhTPS5*, *AhTPS7* and *AhTPS8* genes generally remained unchanged or were repressed by WDS and/ or R (Table 2B). *AhTPS10*, was also induced exclusively in Acru and Acau, whereas it was repressed by WDS in Ahyb. *AhTPS9* was induced by SWDS and R in all species tested with the exception of Ahyb, whereas *AhTPS11* was again induced universally by SWDS, but at lower levels. However, its expression in other conditions tested was more sporadic than in leaves. Importantly, the expression levels of *AhTPS9* during SWDS was significantly higher in roots of Ahypo and Acru, in concordance with their superior WDS tolerance. Also noticeable, was the widespread induction of *AhTPS9-AhTPS11* in Acau. Besides, the expression of all *AhTPP* genes tested was repressed or unaltered by WDS in roots (Table 2C), whereas *AhTRE* ceased to be induced in roots of Ahypo, similarly to the other species examined (Table 2D).

**Table 2.** Relative expression values^1^ of genes involved in trehalose synthesis and degradation in roots of four *Amaranthus* species subjected to two levels of water-deficit stress (moderate [M] and severe [S]) and to subsequent recovery ([R]). Induced (normalized expression values ≥ 2.0; in normal text) and repressed (normalized expression values ≤ 0.5; in italicized text) expression values are emphasized in bold.

| Gene | *A. hypochondriacus* |  |  | *A. cruentus* |  |  | *A. caudatus* |  |  | *A. hybridus* |  |  |
| --- | --- | --- | --- | --- | --- | --- | --- | --- | --- | --- | --- | --- |
|  | M | S | R | M | S | R | M | S | R | M | S | R |
| *AhTPS1* | 0.648 | 0.783 | **0.259** | **0.388** | 1.162 | **0.314** | 0.653 | 0.639 | 1.013 | 1.272 | 1.265 | 0.565 |
| *AhTPS5* | 1.463 | 0.523 | 1.314 | 0.833 | 0.684 | 0.922 | 0.911 | 0.775 | **1.615** | **0.423** | **0.294** | 1.473 |
| *AhTPS7* | **0.470** | 0.935 | **2.427** | **0.481** | 0.977 | 1.440 | 0.621 | 0.635 | 1.444 | 0.817 | 0.668 | 0.841 |
| *AhTPS8* | **1.715** | 0.715 | 0.939 | 0.713 | 0.991 | 0.760 | **1.877** | 1.114 | 1.478 | **0.384** | **0.285** | 0.616 |
| *AhTPS9* | 1.371 | **2.206** | **1.549** | 1.004 | **3.402** | **1.535** | 1.392 | **1.836** | **1.634** | **0.397** | **1.706** | **0.466** |
| *AhTPS10* | 0.536 | 0.810 | 0.695 | 1.154 | **2.562** | 0.868 | 1.321 | **1.512** | **2.212** | **0.325** | **0.356** | 0.888 |
| *AhTPS11* | 1.189 | **6.624** | 1.202 | 0.662 | **8.619** | **2.289** | **1.631** | **4.945** | **3.003** | 0.614 | **3.490** | 0.570 |
| *AhTPPA* | 0.920 | 0.705 | **0.250** | 0.640 | 0.850 | 0.706 | 0.703 | 1.108 | 1.062 | 0.631 | **0.314** | 0.722 |
| *AhTPPD* | 1.452 | **0.396** | **0.197** | 1.188 | 0.750 | 0.776 | 1.035 | 1.329 | 0.744 | **0.460** | **0.256** | 0.594 |
| *AhTPPI* | 0.804 | 0.786 | 0.619 | 0.858 | 1.133 | **0.469** | 0.968 | **0.441** | 0.967 | **0.310** | **0.253** | 0.529 |
| *AhTRE* | 0.854 | 0.711 | 0.722 | 1.192 | 1.257 | 0.615 | 0.526 | 0.622 | 1.346 | 0.520 | **0.495** | 1.058 |

<sup>1</sup>Calculated according to the comparative cycle threshold method (Livak and Schmittgen, 2001) using the *AhACT7*, *AhEF1a* and *AhβTub5* amaranth genes for data normalization.

Tre levels were measured by GC-MS, considered more accurate than HPAEC (Quéro *et al.*, 2013). The results show that a 2-to-3-fold Tre accumulation was induced by both MWDS, SWDS, and sometimes in R, in both leaves (Fig. 1A) and roots (Fig. 1B) of all amaranth species. However, the effect was more noticeable in roots and was stronger in WDS-susceptible species. Tre contents showed a poor correlation with the expression of Tre biosynthesis-related genes. This lack of synchronicity suggested that its accumulation may have involved a post-translational activation of class I TPS1 enzyme(s) (Delorge *et al.*, 2015), an event that remains poorly understood (Rubio-Texeira *et al.*, 2016). Tre accumulation could have also reflected the weak induction of Tre catabolism genes observed, similar to related studies that connected Tre accumulation with TRE inactivation (Goddijn *et al.*, 1997; Müller *et al.*, 2001).

**Figure 1.**
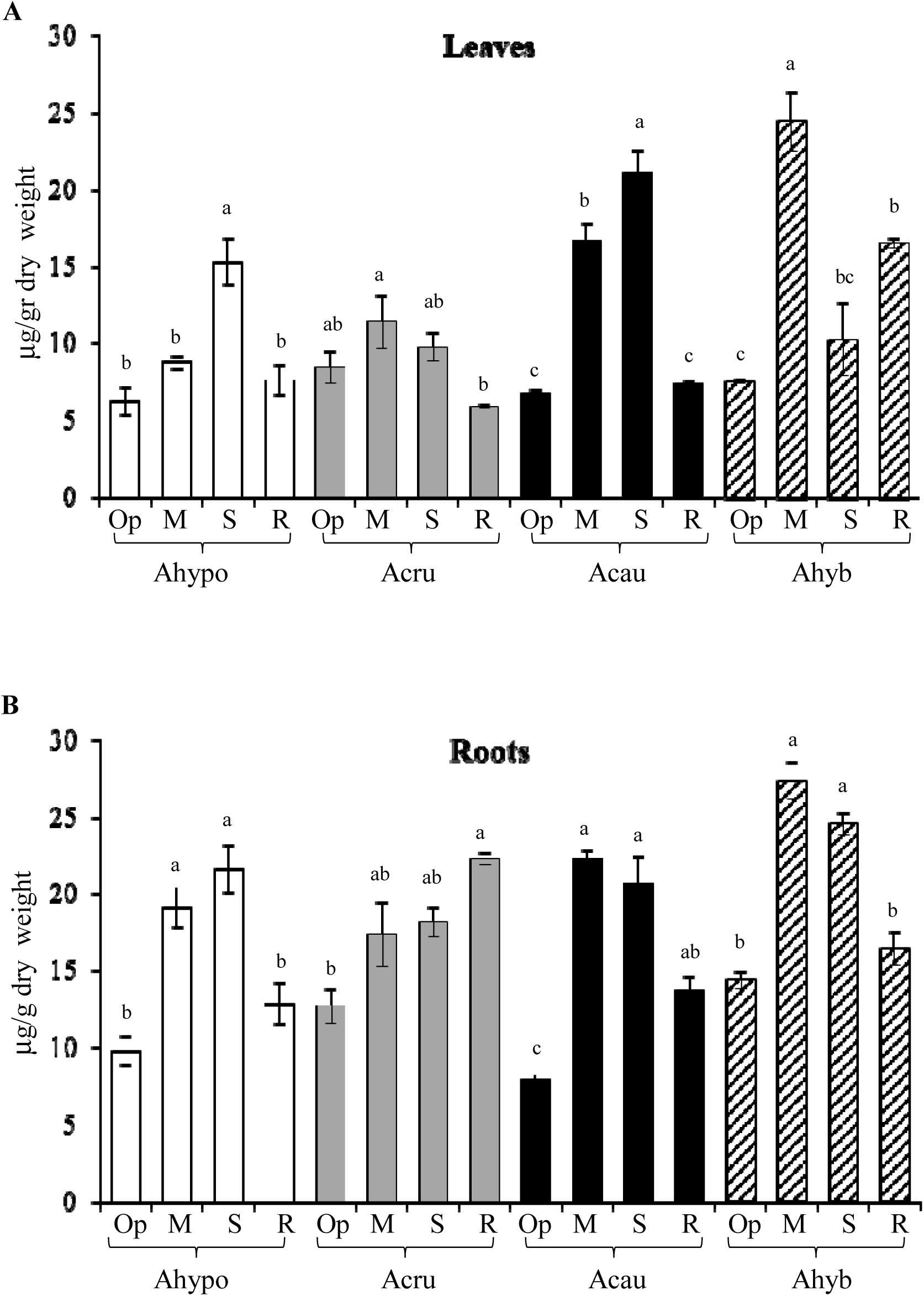
Trehalose content was quantified by GC/ MS in (**A**) leaf and (**B**) root extracts of four species of amaranth plants (i. e., *Amaranthus hypochondriacus* [Ahypo], *A. cruentus* [Acru], *A. caudatus* [Acau] and *A. hybridus* [Ahyb]) growing in optimal conditions (Op), subjected to moderate or severe water deficit stress (MWDS and SWDS, respectively) or allowed to recover from SWDS by restoring watering for 1 day (R).or severe and 1 day after normal watering was restored (R). Different letters over the bars represent statistically significant differences at *P* ≤ 0.05 (Tukey Kramer test). Bars and error bars indicate mean values and ES, respectively (n = 3 pools of four plants each). The results shown are those obtained from a representative experiment that was repeated in the spring-summer and summer-autumn seasons of 2014, respectively, with similar results.

WDS and R induced the expression *of AhSnRAK* in leaves of WDS-susceptible plants and in roots of WDS-tolerant species. *AhSnRK1*a expression remained practically unchanged except for sporadic down- or up-regulated events in leaves and roots. Conversely, *AhSnRK2.1* and *AhSnRK2.2* were negatively affected in leaves of WDS-tolerant plants but induced by WDS in AHyb. Their expression remained unaffected in roots (Supplementary Tables S2, S3).

Regarding RFO-biosynthesis genes, *AhGolS1* and *AhRafS* were almost universally induced by WDS in amaranth leaves, whose expression tended to be highest during SWDS (Table 3). The induction of these genes was lower or returned to basal levels, in R. On the other hand, foliar expression of *AhGolS2* was mostly unaffected by WDS. The expression of *AhStaS*, invariable in leaves of Ahypo and Ahyb, was intermittent in Acau and Acru.

**Table 3.**
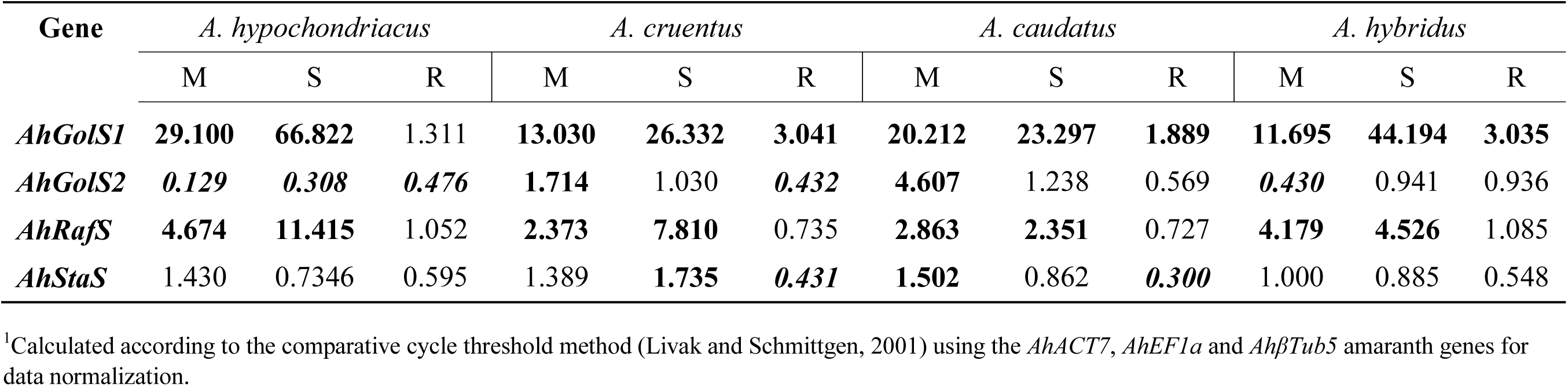
Relative expression values^1^ of genes involved in the biosynthesis of raffinose family oligosaccharides in leaves of four *Amaranthus* species subjected to two levels of water-deficit stress (moderate [M] and severe [S]) and to subsequent recovery ([R]). Induced (normalized expression values ≥ 2.0; in normal text) and repressed (normalized expression values ≤ 0.5; in italicized text) expression values are emphasized in bold.

| Gene | *A. hypochondriacus* |  |  | *A. cruentus* |  |  | *A. caudatus* |  |  | *A. hybridus* |  |  |
| --- | --- | --- | --- | --- | --- | --- | --- | --- | --- | --- | --- | --- |
|  | M | S | R | M | S | R | M | S | R | M | S | R |
| *AhGolS1* | **29.100** | **66.822** | 1.311 | **13.030** | **26.332** | **3.041** | **20.212** | **23.297** | **1.889** | **11.695** | **44.194** | **3.035** |
| *AhGolS2* | *0.129* | *0.308* | *0.476* | **1.714** | 1.030 | *0.432* | **4.607** | 1.238 | 0.569 | *0.430* | 0.941 | 0.936 |
| *AhRafS* | **4.674** | **11.415** | 1.052 | **2.373** | **7.810** | 0.735 | **2.863** | **2.351** | 0.727 | **4.179** | **4.526** | 1.085 |
| *AhStaS* | 1.430 | 0.7346 | 0.595 | 1.389 | **1.735** | *0.431* | **1.502** | 0.862 | *0.300* | 1.000 | 0.885 | 0.548 |

<sup>1</sup>Calculated according to the comparative cycle threshold method (Livak and Schmittgen, 2001) using the *AhACT7*, *AhEF1a* and *AhβTub5* amaranth genes for data normalization.

*AhGolS1* expression in roots of WDS-treated amaranth plants was intensely induced by SWDS, notably in Acru (Table 4). *AhGolS1* expression in roots of Ahypo during SWDS was also high, being 1.8- to 3.6-fold higher than those detected in Acau and Ahyb, respectively. Thus, *AhGolS1* expression pattern in roots of grain amaranth plants also coincided with their WDS tolerance. Conversely, *AhGolS2* was almost universally induced in response to WDS in roots. Root *AhGolS2* gene expression patterns in response to WDS were mirrored by those produced by the *AhRafS* and *AhStaS* genes, except for the occasional induction of the latter during R. The ca. 3- to 10-fold higher *AhRafS* expression levels detected in roots of Ahypo and Acru subjected to SWDS, compared to those in Acau and Ahyb, also agreed with their increased WDS tolerance.

**Table 4.** Relative expression values^1^ of genes involved in the biosynthesis of raffinose family oligosaccharides in roots of four *Amaranthus* species subjected to two levels of water-deficit stress (moderate [M] and severe [S]) and to subsequent recovery ([R]). Induced (normalized expression values ≥ 2.0; in normal text) and repressed (normalized expression values ≤ 0.5; in italicized text) expression values are emphasized in bold.

| Gene | *A. hypochondriacus* |  |  | *A. cruentus* |  |  | *A. caudatus* |  |  | *A. hybridus* |  |  |
| --- | --- | --- | --- | --- | --- | --- | --- | --- | --- | --- | --- | --- |
|  | M | S | R | M | S | R | M | S | R | M | S | R |
| *AhGolS1* | **22.202** | **136.744** | 0.819 | **14.745** | **248.857** | **2.128** | **18.018** | **75.932** | 1.088 | **17.009** | **37.691** | **1.870** |
| *AhGolS2* | 0.673 | **3.471** | 0.261 | **2.697** | **3.345** | 1.198 | **2.401** | **2.539** | 1.124 | **2.355** | **2.328** | 0.696 |
| *AhRafS* | **3.426** | **14.981** | 1.270 | **1.795** | **15.190** | 1.008 | **2.268** | **5.426** | 0.539 | **3.699** | 1.407 | 0.510 |
| *AhStaS* | **2.960** | **6.219** | 1.378 | **6.583** | **5.126** | **1.553** | **5.525** | **5.089** | **2.381** | **2.943** | **2.758** | 1.034 |

<sup>1</sup>Calculated according to the comparative cycle threshold method (Livak and Schmittgen, 2001) using the *AhACT7*, *AhEF1a* and *AhβTub5* amaranth genes for data normalization.

The RFO accumulation pattern in leaves and roots (Fig. 2, 3) partially coincided the expression of RFO biosynthesis-related genes (Tables 3, 4). Raf accumulation in leaves could be likewise suggested as another contributing factor to the increased WDS tolerance observed in Ahypo and Acru. In contrast, practically no accumulation of Gol was detected in leaves of all species, irrespective of their treatment, suggesting an active utilization of this precursor for the synthesis of RFOs. On the other hand, MI content remained unaltered in leaves of Ahypo plants (Fig. 2A), but accumulated, particularly during SWDS, in Acru and Acau (Fig. 2B, C). Sta content was minimal in leaves of all species and changes were small and sporadic (Fig. 2A, B, D). Similarly Ver content in Ahypo and Acru (Fig. 2A, B) was modest and static. However, Ver levels increased to ca. 5-fold higher levels than controls in response to WDS in Acau and Ahyp (Fig. 2C, D). The above results suggest that Raf/ Ver ratios in leaves could constitute a marker of WDS tolerance in amaranth.

**Figure 2.**
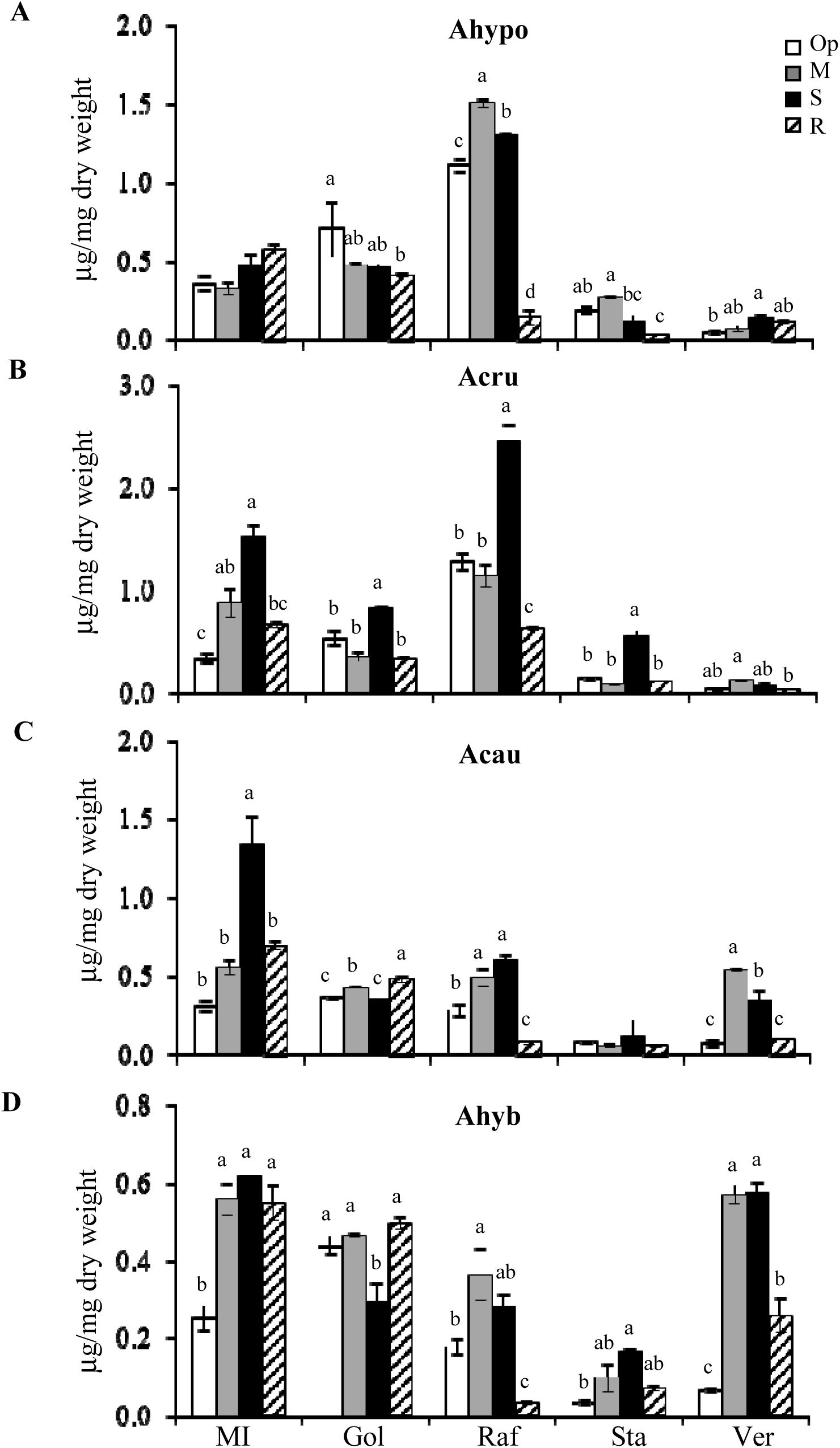
Raffinose family oligosaccharides (RFOs) were quantified by HPAEC-PAD in leaf extracts of four species of amaranth plants: (**A**) *Amaranthus hypochondriacus* [Ahypo], (**B**) *A. cruentus* [Acru], (**C**) *A. caudatus* [Acau], and (**D**) *A. hybridus* [Ahyb]) growing in optimal conditions (Op; empty bars), subjected to moderate (M) or severe (S) water deficit stress (gray and black bars respectively) or allowed to recover from S, 1 day after normal watering was restored (R; striped bars). The RFOs and their respective precursors analyzed were myo-inositol (MI), galactinol (Gol), raffinose (Raf), staquiose (Sta) and verbascose (Ver). Different letters over the bars represent statistically significant differences at *P* ≤ 0.05 (Tukey Kramer test). Bars and error bars indicate mean values and ES, respectively (n = 3 pools of four plants each). The results shown are those obtained from a representative experiment that was repeated in the spring-summer and summer-autumn seasons of 2014, respectively, with similar results.

**Figure 3.**
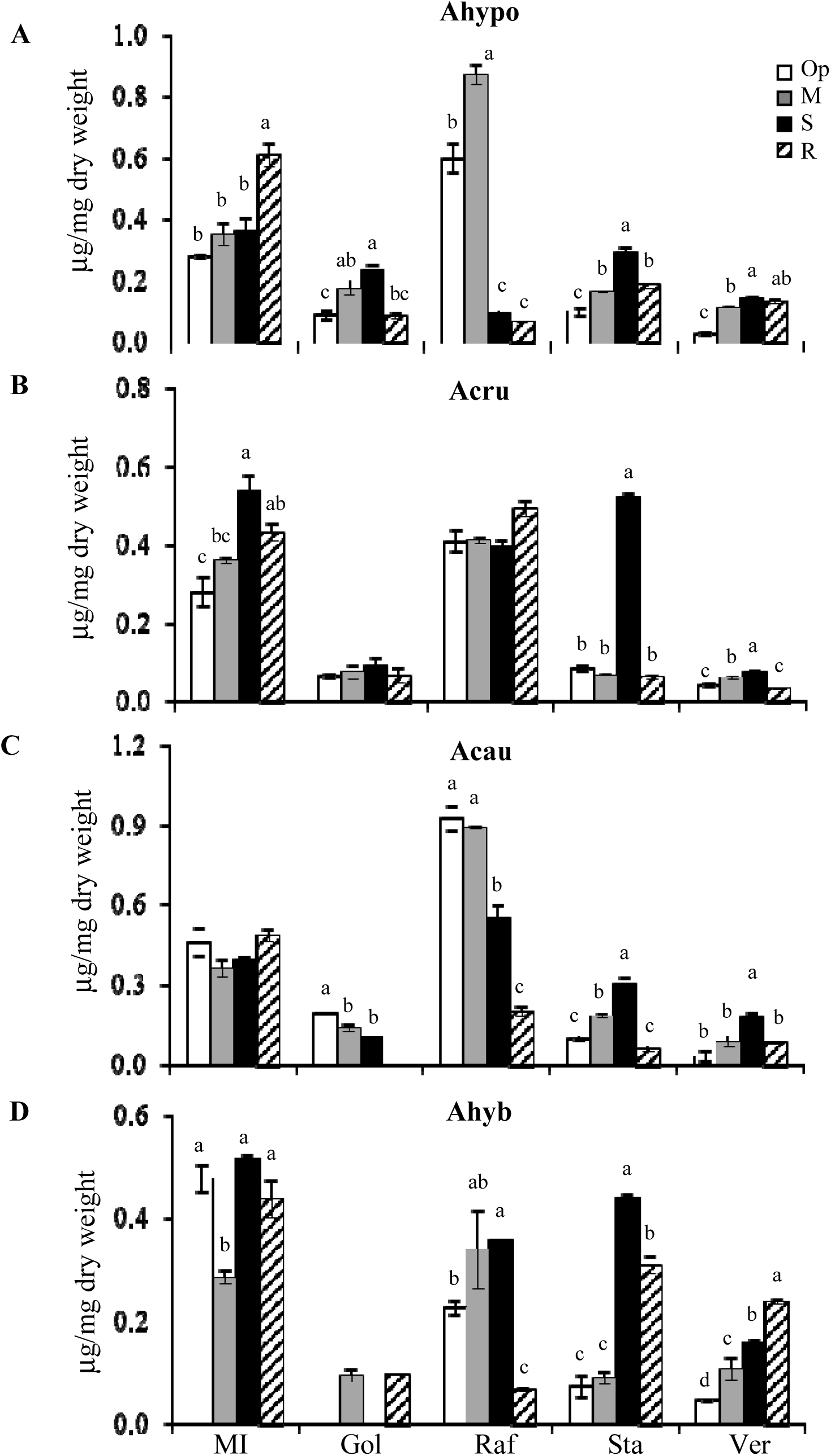
Raffinose family oligosaccharides (RFOs) were quantified by HPAEC-PAD in root extracts of four species of amaranth plants: (**A**) *Amaranthus hypochondriacus* [Ahypo], (**B**) *A. cruentus* [Acru], (**C**) *A. caudatus* [Acau], and (**D**) *A. hybridus* [Ahyb]) growing in optimal conditions (Op; empty bars), subjected to moderate (M) or severe (S) water deficit stress (gray and black bars respectively) or allowed to recover from S, 1 day after normal watering was restored (R; striped bars). The RFOs and their respective precursors analyzed were myo-inositol (MI), galactinol (Gol), raffinose (Raf), staquiose (Sta) and verbascose (Ver). Different letters over the bars represent statistically significant differences at *P* ≤ 0.05 (Tukey Kramer test). Bars and error bars indicate mean values and ES, respectively (n = 3 pools of four plants each). The results shown are those obtained from a representative experiment that was repeated in the spring-summer and summer-autumn seasons of 2014, respectively, with similar results.

The root RFO results differed and had a lower correspondence with WDS tolerance in amaranth. Raf did not to accumulate in response to WDS in Ahypo and Acru (Fig. 3A, B). In Ahyb, Raf content fluctuations in roots were similarly erratic than those in leaves (Fig. 3D), whereas the ca. 2-fold higher basal Raf content in roots of Acau was drastically reduced by SWDS and R, similarly to Ahypo (Fig. 3C). SWDS conditions also induced the accumulation of MI in roots of Acru (Fig. 3B), while a significant increase occurred in Ahypo roots during R (Fig. 3A). Basal Gol contents in roots were ca. 2-fold lower than in leaves, and undetectable under certain conditions in roots of Ahyb (Fig. 3D). Sta contents remained low in roots and also showed a tendency to accumulate in response to SWDS. Contrary to leaves, root Sta accumulation was significantly increased by SWDS in all species tested (Fig. 3C). Likewise, Ver content increased in roots of all species in response to SWDS (Fig. 3A-D). Curiously, Ahypo and Ahyb accumulated almost identical Ver contents in response to WDS o R treatments (Fig. 3A, D).

The significantly higher foliar accumulation of Raf in Ahypo and Acru observed in response to MWDS and SWDS correlated with significantly augmented *AhGolS1* and *AhRafS* expression levels (Table 3). In roots this association was not found, although these genes were expressed to ca. 10-fold higher levels than those detected in leaves under similar conditions (Table 4). The reason(s) why the intense induction of these genes did not translate into high contents of Raf and perhaps other RFOs in roots remains unknown. Contrarily, changes in *AhStaS* expression in response to WDS and R agreed with root Sta levels (Table 4; Fig. 3). However, this correspondence was not detected in leaves (Table 3; Fig. 2). The lack of coincidence between RFO content and their correspondent gene expression in some amaranth species could be explained by the possibility that these were being converted to putatively larger RFO, whose structure is yet to be determined. In this respect, several unknown compounds having longer retention times (RTs), and perhaps larger sizes, were detected (Supplementary Fig. S4-S7). Peaks with RTs of 16.1, 22.2 and 33.8 min were abundant in leaves WDS tolerant Ahypo and Acru, particularly in the former. Thus, they could be considered as contributors to WDS tolerance in these species. Contrarily, two peaks with RTs of ca. 16.8 and 21.1 min accumulated in roots of most treated plants, noticeably during SWDS and R. Curiously, both compounds were more abundant in WDS susceptible species. Thin-layer chromatography traces of both leaf and root crude extracts (Supplementary Fig. S8) show bands having differential intensity that could correspond to these unknown compounds, whose nature remains to be determined experimentally.

WDS marker genes were not uniformly expressed in treated plants; they varied depending on the treatment applied, organ examined and species. In leaves, *AhABI5* and *AhLEA14* were the only genes induced almost uniformly across species by WDS (Table 5A), although *AhLEA14* expression was several-fold higher than *AhABI5*, and was induced in all conditions tested. Conversely, *AhRAB18* was sporadically induced by WDS in Acau and Ahyb, whereas it remained practically unchanged in Ahypo and Acru. Contrariwise, *AhDREB2C* was induced by all treatments in Ahypo only. All marker genes were more intensely induced in roots (Table 5B), distinctly in Ahypo, Acru and Acau, whereas they remained mostly unaltered in Ahyb. Importantly, marker genes reached their highest expression in both leaves and roots of treated Ahypo plants, in correspondence with their superior WDS tolerance.

**Table 5.** Relative expression values^1^ of abscisic acid (ABA) marker genes in leaves of four *Amaranthus* species subjected to two levels of water-deficit stress (moderate [**M**] and severe [**S**]) and to subsequent recovery ([**R**]). Induced (normalized expression values ≥ 2.0; in normal text) and repressed (normalized expression values ≤ 0.5; in italicized text) expression values are emphasized in bold.

| Gene | *A. hypochondriacus* |  |  | *A. cruentus* |  |  | *A. caudatus* |  |  | *A. hybridus* |  |  |
| --- | --- | --- | --- | --- | --- | --- | --- | --- | --- | --- | --- | --- |
|  | M | S | R | M | S | R | M | S | R | M | S | R |
| *AhABI5* | **2.005** | **3.047** | **2.202** | **1.937** | **4.014** | 0.726 | **3.244** | 1.469 | 1.317 | **2.437** | **2.590** | **2.507** |
| *AhDREB* | **5.300** | **8.754** | **3.867** | 0.858 | 1.251 | 0.631 | **1.655** | 0.921 | 0.739 | 0.665 | **1.689** | 1.159 |
| *AhRAB18* | 0.776 | 0.789 | 0.963 | 0.885 | 0.843 | *0.350* | **1.642** | 1.233 | 0.861 | 0.502 | **1.815** | 1.052 |
| *AhLEA14* | **21.284** | **64.530** | **5.835** | **16.518** | **31.398** | **2.136** | **49.051** | **120.856** | **4.553** | **16.745** | **46.504** | **2.539** |

<sup>1</sup>Calculated according to the comparative cycle threshold method (Livak and Schmittgen, 2001) using the *AhACT7*, *AhEF1a* and *AhβTub5* amaranth genes for data normalization.

Pro levels were significantly higher in leaves of WDS-tolerant species, where the highest Pro contents accumulated in response to SWDS (Fig. 4A). Contrariwise, Pro accumulation in roots (Fig. 4B), did not vary much between amaranth species, where a significant increase was only observed in SWDS (although ca. 2.5-fold lower than in leaves). Significantly higher root Pro levels were also detected in MWDS and R in Ahyb, whereas the lowest Pro accumulation occurred in Acau. In contrast, NSCs content fluctuations in leaves and roots were consistent with the contrasting WDS tolerance observed between amaranth species. Thus, all NSCs were significantly lower in leaves of WDS-tolerant species, distinctly during SWDS (Figs. 5). In roots (Fig. 6), the NSC content variation in Ahypo was manifestly different. Thus Glu and Fru were the highest detected and Suc and starch levels the lowest. This occurred independently of the treatment analyzed. Lower hexose (Hex) content in leaves of Ahypo was in agreement with the WDS-unresponsive invertase activity observed (Fig. 7A-C), whereas consistently higher Glu and Fru contents in leaves of treated Acau and Ahyb plants coincided with increased cell wall invertase (CWI) (in most conditions tested; Fig. 7A), and to augmented vacuolar (VI) and cytoplasmic invertase activities (CI), mostly during R (Fig. 7A-C). In Acru, a gradual increase in Glu and Fru observed during WDS and R, could be attributed to an increased activity in all three invertases tested (Fig. 7A-C), mostly during SWDS. Nevertheless, its foliar Hex levels tended to be the lowest, together with Ahypo. Also intriguing was the Suc peak produced during SWDS in the latter species (Fig. 5A-C).

**Figure 4.**
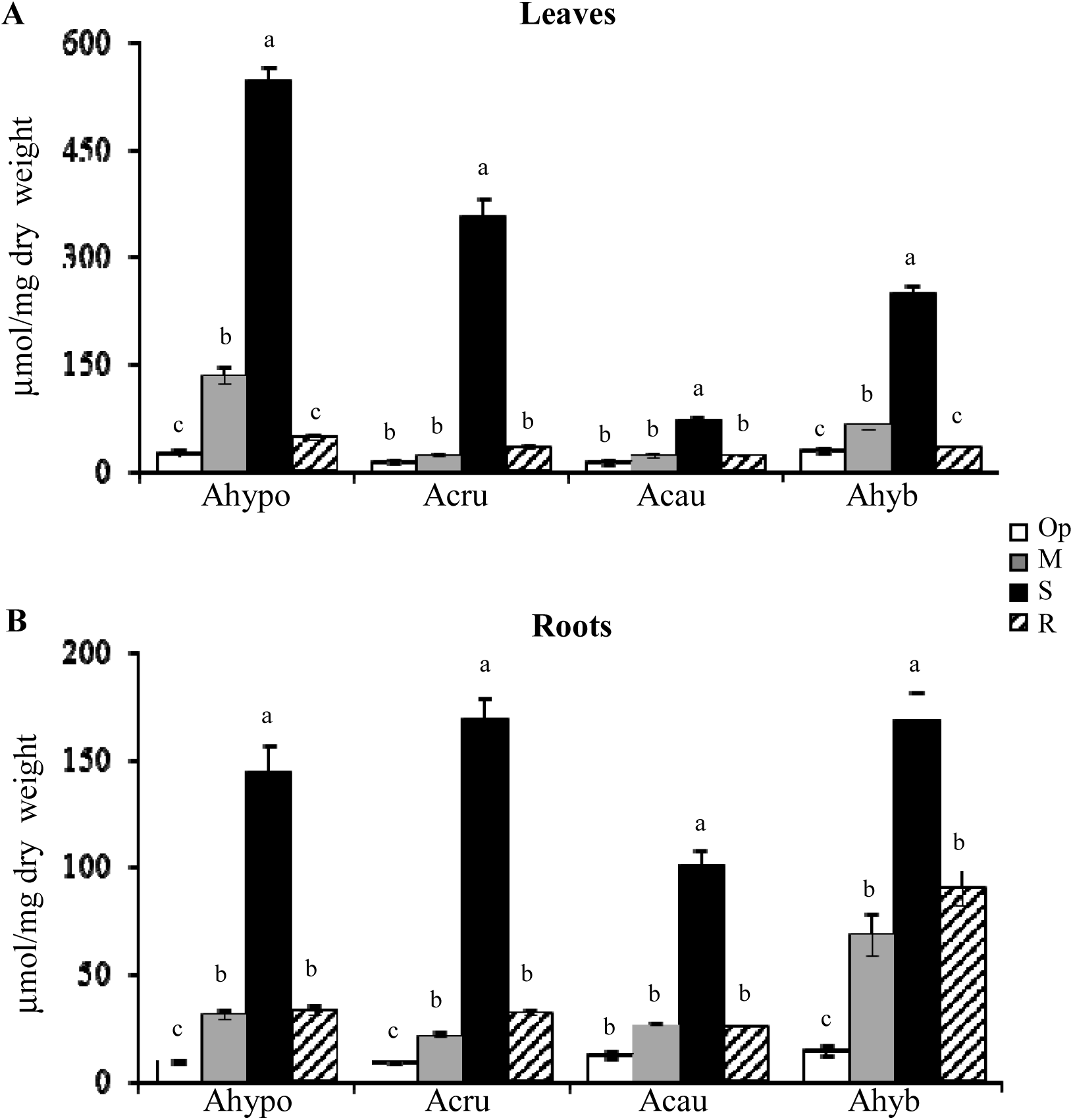
Proline content quantified *in vitro* in (**A**) leaf and (**B**) root extracts of four species of amaranth plants (i. e., *Amaranthus hypochondriacus* [Ahypo], *A. cruentus* [Acru], *A. caudatus* [Acau] and *A. hybridus* [Ahyb]) growing in optimal conditions (Op; empty bars), subjected to moderate (M) or severe (S) water deficit stress (gray and black bars respectively) or allowed to recover from S, 1 day after normal watering was restored (R; striped bars). Different letters over the bars represent statistically significant differences at *P* ≤ 0.05 (Tukey Kramer test). Bars and error bars indicate mean values and ES, respectively (n = 3 pools of four plants each). The results shown are those obtained from a representative experiment that was repeated in the spring-summer and summer-autumn seasons of 2014, respectively, with similar results.

**Figure 5.**
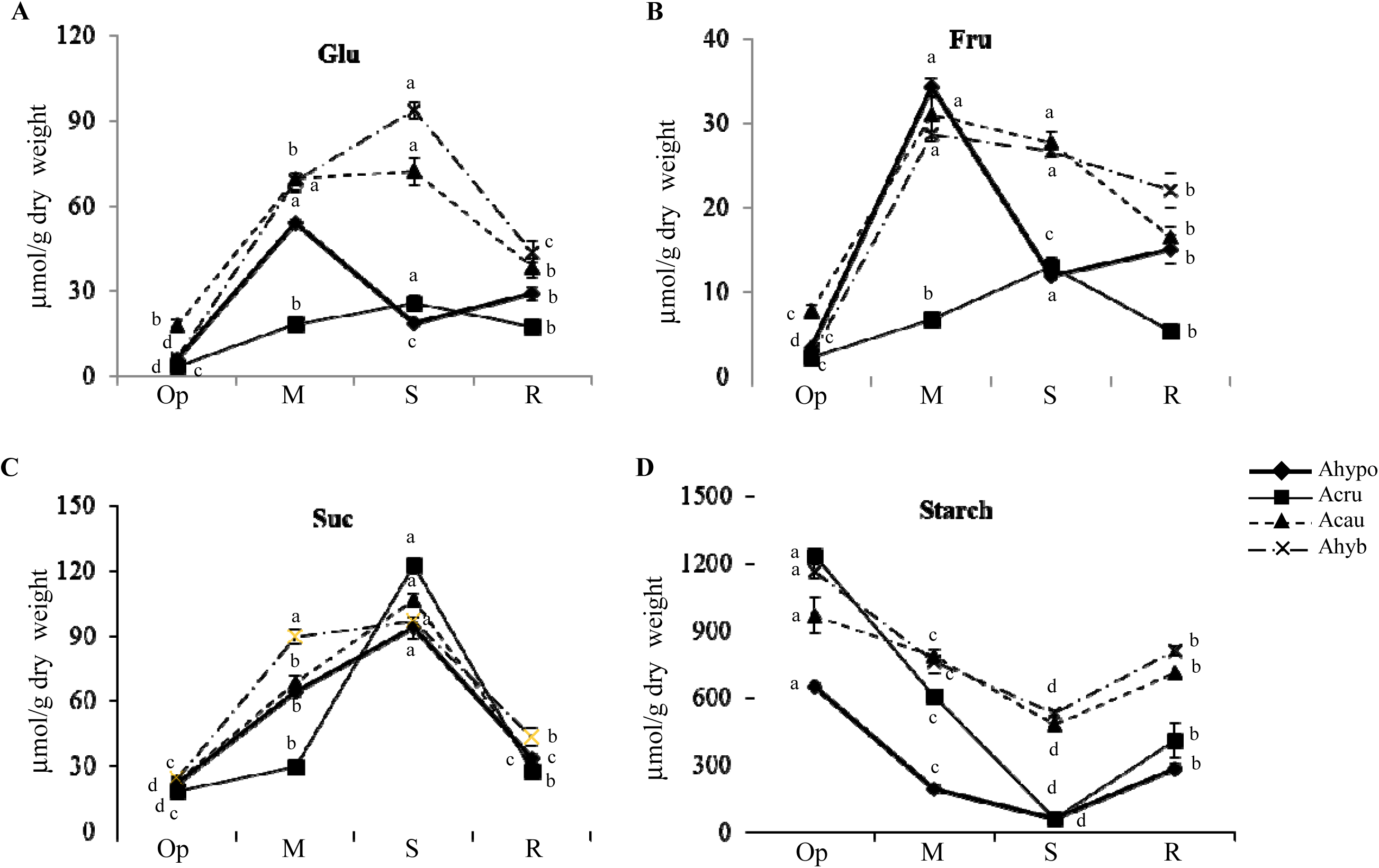
Non-structural carbohydrates (Glucose [Glu], Fructose [Fru], Sucrose [Suc] and starch) content quantified *in vitro* in leaves of four species of amaranth plants (i. e., Amaranthus hypochondriacus [Ahypo; thick continuous line], A. cruentus [Acru; thin continuous line], A. caudatus [Acau; short dash line] and A. hybridus [Ahyb; long chain line]) growing in optimal conditions (Op), subjected to moderate (M) or severe (S) water deficit stress, or allowed to recover from S, 1 day after normal watering was restored (R). Different letters over the lines represent statistically significant differences at *P* ≤ 0.05 (Tukey Kramer test). Bars and error bars indicate mean values and ES, respectively (n = 3 pools of four plants each). The results shown are those obtained from a representative experiment that was repeated in the spring-summer and summer-autumn seasons of 2014, respectively, with similar results.

**Figure 6.**
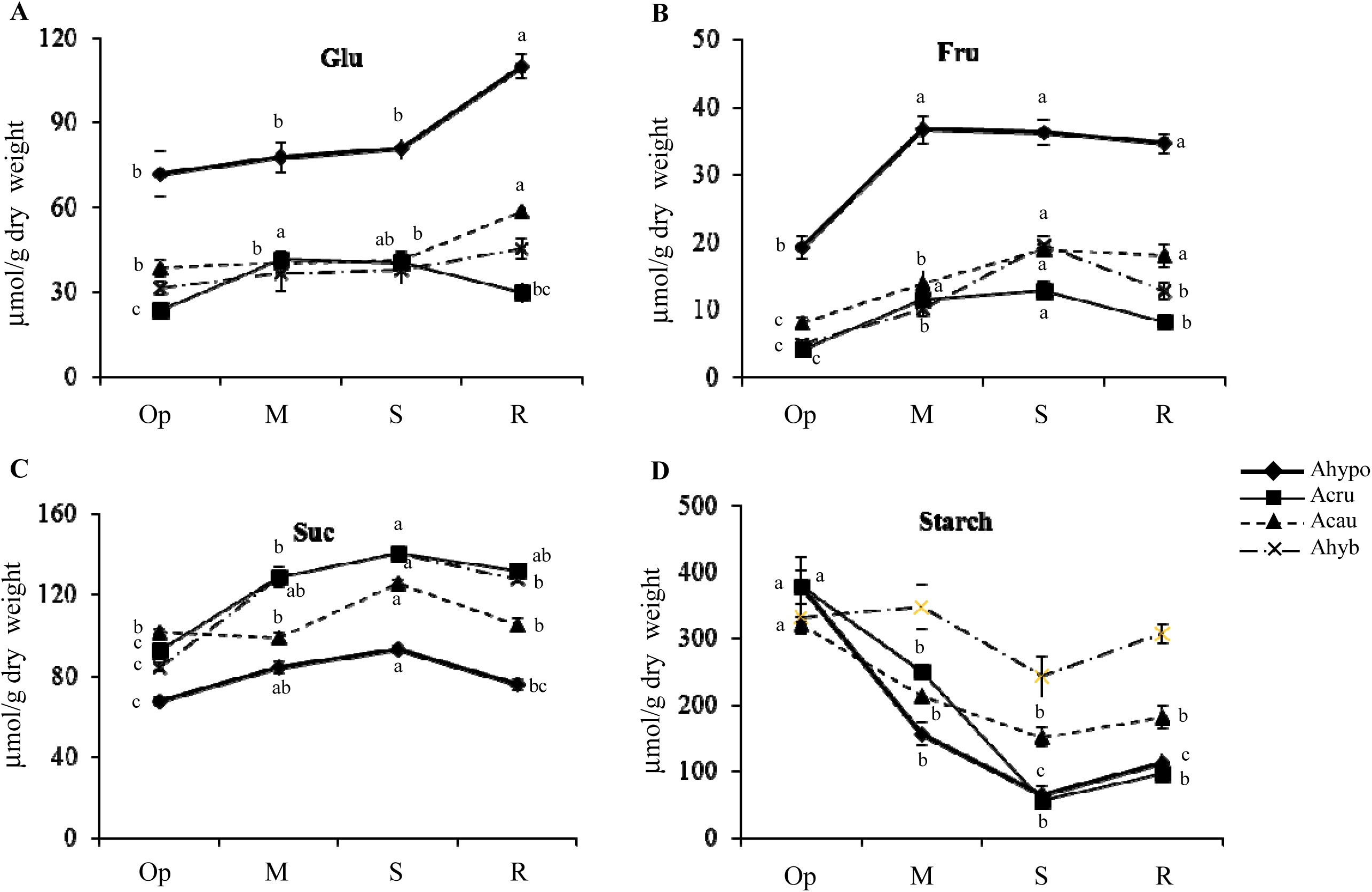
Non-structural carbohydrates (Glucose [Glu], Fructose [Fru], Sucrose [Suc] and starch) content quantified *in vitro* in roots of four species of amaranth plants (i. e., Amaranthus hypochondriacus [Ahypo; thick continuous line], A. cruentus [Acru; thin continuous line], A. caudatus [Acau; short dash line] and A. hybridus [Ahyb; long chain line]) growing in optimal conditions (Op), subjected to moderate (M) or severe (S) water deficit stress, or allowed to recover from S, 1 day after normal watering was restored (R). Different letters over the lines represent statistically significant differences at *P* ≤ 0.05 (Tukey Kramer test). Bars and error bars indicate mean values and ES, respectively (n = 3 pools of four plants each). The results shown are those obtained from a representative experiment that was repeated in the spring-summer and summer-autumn seasons of 2014, respectively, with similar results.

**Figure 7.**
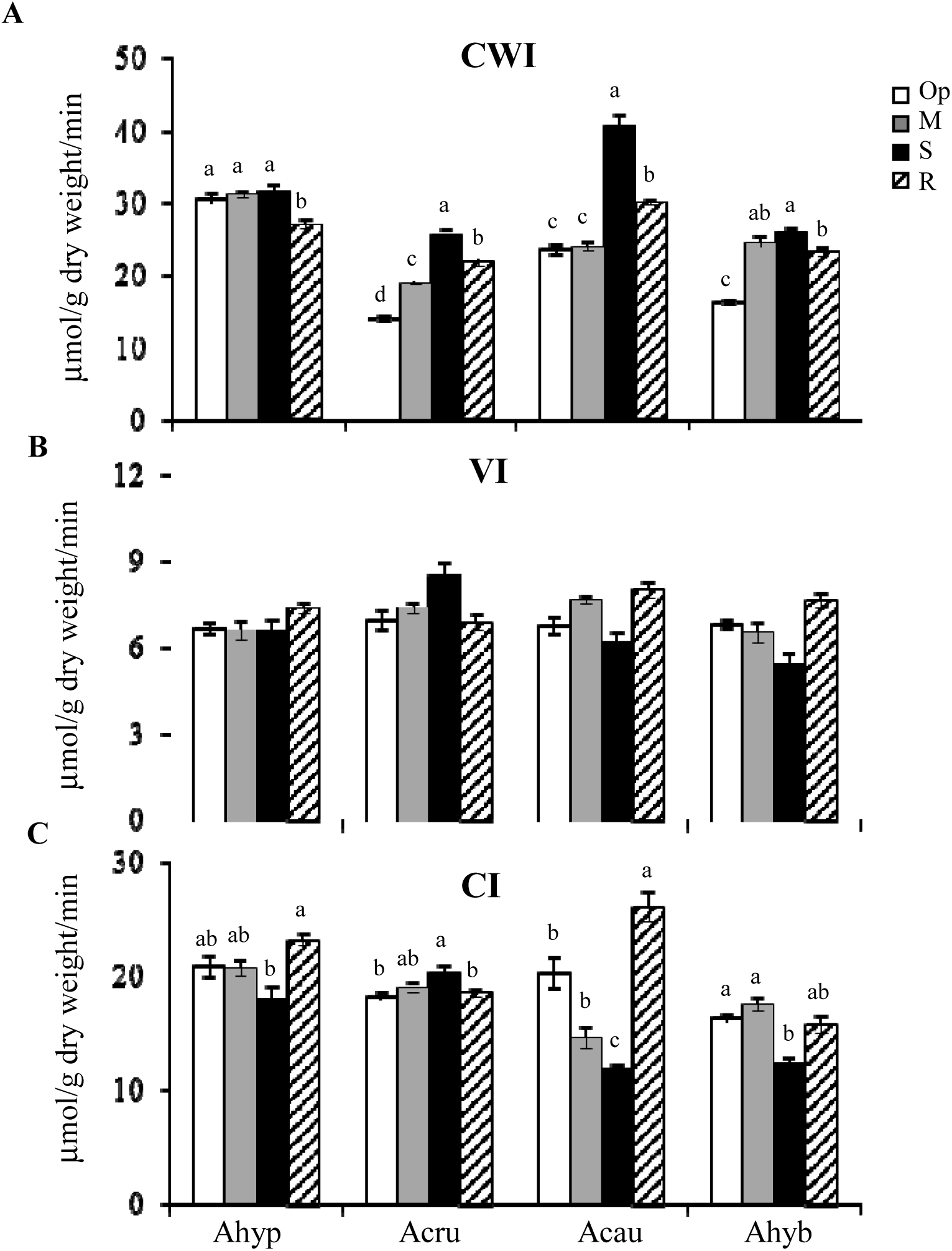
(**A**) Cell wall invertase (CWI), (**B**) vacuolar invertase VI), and (**C)** neutral cytoplasmic invertase (CI) activities determined *in vitro* in leaf extracts of four species of amaranth plants: *Amaranthus hypochondriacus* [Ahypo], *A. cruentus* [Acru], (C), *A. caudatus* [Acau], and (D) *A. hybridus* [Ahyb], growing in optimal conditions (Op; empty bars), subjected to moderate (M) or severe (S) water deficit stress (gray and black bars respectively) or allowed to recover from S, 1 day after normal watering was restored (R; striped bars). Different letters over the bars represent statistically significant differences at *P* ≤ 0.05 (Tukey Kramer test). Bars and error bars indicate mean values and ES, respectively (n = 3 pools of four plants each). The results shown are those obtained from a representative experiment that was repeated in the spring-summer and summer-autumn seasons of 2014, respectively, with similar results.

Conversely, the high Hex/ Suc ratio observed in roots of treated Ahypo plants was consistent with increased CWI and VI activity (Fig. 8A, B), and with a strong induction of SuSy activity in SWDS (Fig. 9). Lower SuSy activities, combined with repressed and/ or unchanged invertase activity in roots of Acau and Ahyb treated plants were consistent with their lower Hex/ Suc ratios. No SuSy activity was detected in leaves.

**Figure 8.**
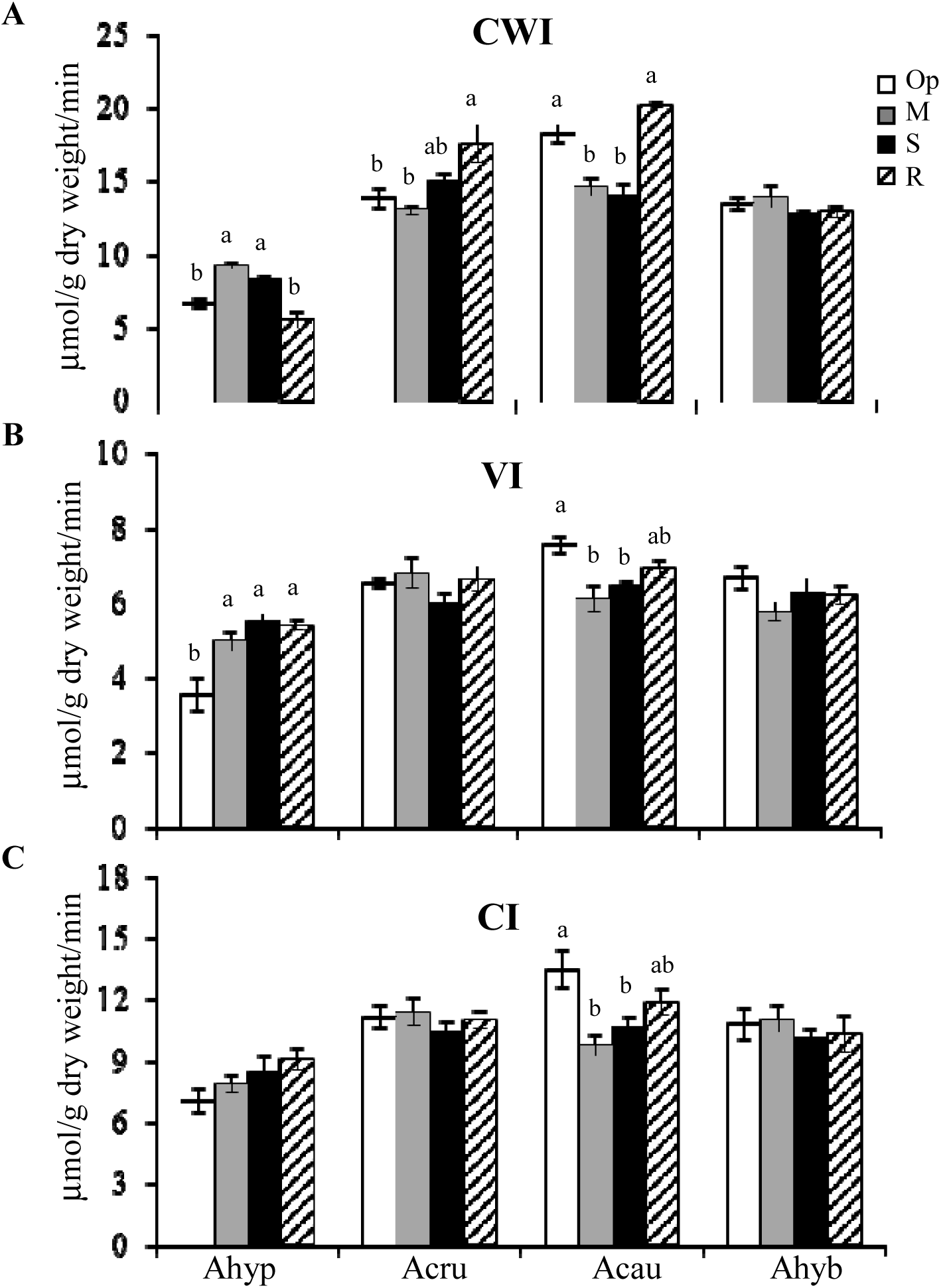
(**A**) Cell wall invertase, (**B**) vacuolar invertase, and (**C)** neutral cytoplasmic invertase activities determined *in vitro* in root extracts of four species of amaranth plants: *Amaranthus hypochondriacus* (Ahypo), *A. cruentus* (Acru), *A. caudatus* (Acau), and *A. hybridus* (Ahyb), growing in optimal conditions (Op; empty bars), subjected to moderate (M) or severe (S) water deficit stress (gray and black bars respectively) or allowed to recover from S, 1 day after normal watering was restored (R; striped bars). Different letters over the bars represent statistically significant differences at *P* ≤ 0.05 (Tukey Kramer test). Bars and error bars indicate mean values and ES, respectively (n = 3 pools of four plants each). The results shown are those obtained from a representative experiment that was repeated in the spring-summer and summer-autumn seasons of 2014, respectively, with similar results.

**Figure 9.**
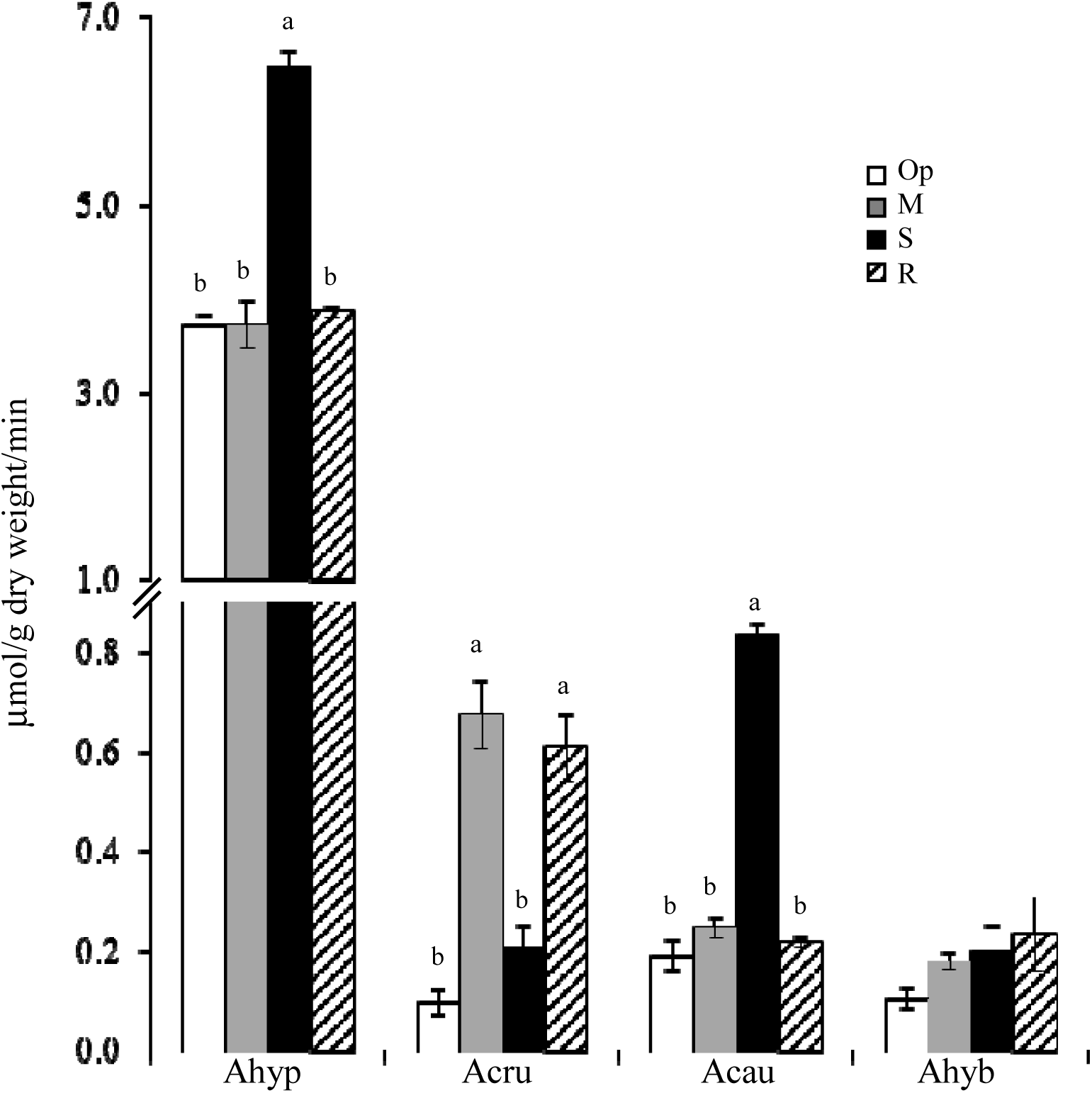
Sucrose synthase determined *in vitro* in root extracts of four species of amaranth plants: *Amaranthus hypochondriacus* (Ahypo), *A. cruentus* (Acru), *A. caudatus* (Acau), and *A. hybridus* (Ahyb), growing in optimal conditions (Op; empty bars), subjected to moderate (M) or severe (S) water deficit stress (gray and black bars respectively) or allowed to recover from S, 1 day after normal watering was restored (R; striped bars). Different letters over the bars represent statistically significant differences at *P* ≤ 0.05 (Tukey Kramer test). Bars and error bars indicate mean values and ES, respectively (n = 3 pools of four plants each). The results shown are those obtained from a representative experiment that was repeated in the spring-summer and summer-autumn seasons of 2014, respectively, with similar results.

The above results indicated WDS had a different effect on the NSC content of leaves and roots in tolerant Ahypo and Acru, compared to susceptible Acau and Ahyb. Thus, leaves of tolerant amaranths tended to have lower Glu, Fru, and starch contents. The effect was drastic during SWDS, particularly for starch reserves, which were almost depleted (Fig. 5D, 6D). In contrast, constitutive Glu levels in roots of Ahypo plants were significantly higher than those in all others and increased significantly in R (Fig. 5A), whereas constitutively high Fru levels, further increased after WDS treatment (Fig. 5B). Amylolytic activity was almost uniformly induced in leaves of all treated plants (Fig. 10A), whereas is induction by all treatments was observed only in roots of Ahypo (Fig. 10B). This contrast suggests that additional starch degradation mechanisms contributed to the starch depletion observed in leaves and roots of WDS-tolerant amaranths (Grennan, 2006; Turesson *et al.*, 2014).

**Figure 10.**
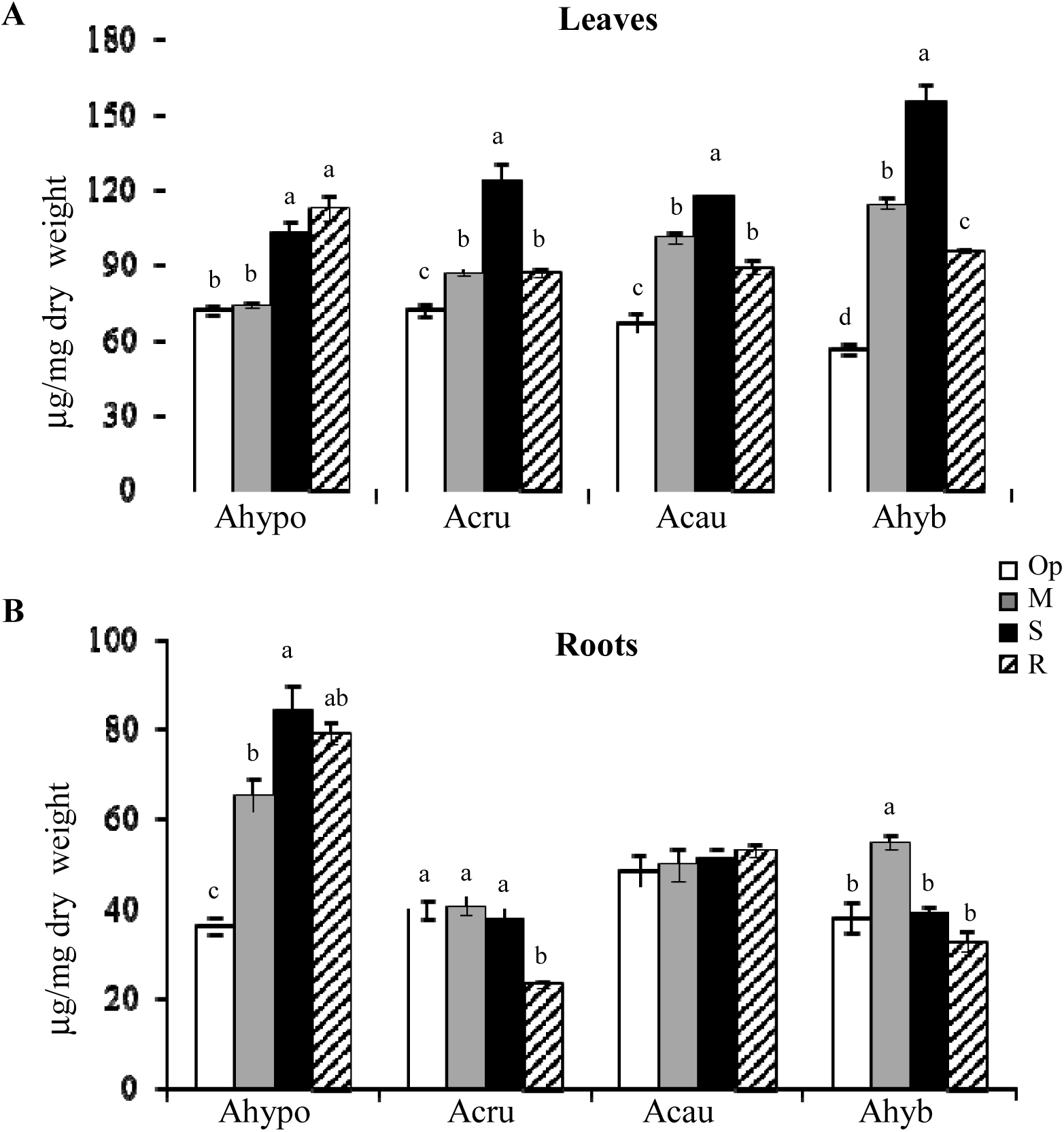
Amylase activity quantified *in vitro* in (**A**) leaf and (**B**) root extracts of four species of amaranth plants (i. e., *Amaranthus hypochondriacus* [Ahypo], *A. cruentus* [Acru], *A. caudatus* [Acau] and *A. hybridus* [Ahyb]) growing in optimal conditions (Op; empty bars), subjected to moderate (M) or severe (S) water deficit stress (gray and black bars respectively) or allowed to recover from S, 1 day after normal watering was restored (R; striped bars). Different letters over the bars represent statistically significant differences at *P* ≤ 0.05 (Tukey Kramer test). Bars and error bars indicate mean values and ES, respectively (n = 3 pools of four plants each). The results shown are those obtained from a representative experiment that was repeated in the spring-summer and summer-autumn seasons of 2014, respectively, with similar results.

## 4. Discussion

It was previously shown that the WDS response in Ahypo roots included the accumulation of osmolytes and increased levels of ROS scavenging and heat shock proteins, together with the induction of certain TFs (Huerta-Ocampo *et al.*, 2011). Several other amaranth genes have been subsequently proposed as possible contributing factors to increased tolerance against several (a)abiotic stresses in grain amaranth, including an orphan gene (Massange-Sánchez *et al.*, 2015), a gene with an unknown function domain (Palmeros-Súarez *et al.*, 2017) and various TF genes (Palmeros Súarez *et al.*, 2015; Massange-Sánchez *et al.*, 2016). The above genes were induced in grain amaranth by several stress conditions and frequently conferred stress tolerance when overexpressed in Arabidopsis plants.

The present study found, however, that WDS tolerance in grain amaranth varied within and between species. Ahypo and Acru tended to be tolerant, whereas Acau, an incompletely domesticated grain amaranth species (Stetter *et al.*, 2017), and Ahyb, an undomesticated species presumed to be their ancestor (Stetter and Schmid, 2017), were susceptible. A battery of molecular and biochemical tests were employed to identify the bases of such difference. Changes in Tre and RSOs accumulation, as well as in the expression of related genes, together with modifications in C mobilization and in Pro content during WDS and in R were monitored. The general unresponsiveness of *AhTPS1* and downstream targets (i.e., *AhSnRK1*) (Tables 1, 2; Supplementary Tables S2, S3) to WDS suggest that the role of T6P-related signaling was probably not a defining factor of WDS tolerance in grain amaranth. A similar prediction could be proposed for Tre (Fig. 1). This was partly in agreement with a study showing that Tre did not protect yeast cells from desiccation (Petitjean *et al.*, 2015) and with others that found no link between increased Tre accumulation and stress tolerance. It was contradictory, however, to evidence connecting Tre accumulation with WDS tolerance (Figueroa and Lunn, 2016). Moreover, it may be suggested that increased foliar Tre levels, could have contributed to WDS susceptibility in Acau and Ahyb, similar to Arabidopsis *tre* null mutants that had increased Tre levels and were more sensitive to drought than WT plants (Van Houtte *et al.*, 2013). Such effect was ascribed to a proposed link connecting Tre metabolism, stomatal conductance and variations in stomata’s responsiveness to ABA.

Moreover, gene expression assays established a poor correlation between other Tre-related genes and increased WDS tolerance in Ahypo, except for a few exceptions: i) the general downregulation of foliar *AhTPS5*, of several other *class II TPS* genes during R, and of the *AhTPPD* and *AhTPPI* genes during WDS; ii) the high expression of *AhTPS9* and *AhTPS11* observed in leaves and roots (together with Acru) during SWDS, and iii) the induction of *AhTRE* during WDS (Tables 1, 2). Past studies have shown that class II TPS proteins have a differential sensitivity to Suc levels in plants (Schluepmann and Paul, 2009), which is important in the context of modified NSCs content observed in response to WDS in amaranth and other plants (Pinheiro and Chaves 2010). This property could explain the increased induction of the *AhTPS9* and *AhTPS11* in WDS-tolerant amaranth. However, the role of these genes in WDS amelioration remains to be determined. The above results were also consistent with the upregulation of *TPS11* and *TRE* detected in stomatal guard cells of sucrose-treated Arabidopsis (Bates *et al.*, 2012). Such coordinated effect was proposed to establish a connection between Tre metabolism, carbohydrate metabolism regulation, and stomatal movements via sugar sensing, and further supported the role of Tre in the regulation of stomatal behavior.

The significantly higher upregulation of *AhTPS9* and *AhTPS11* under SWDS in sucrose-depleted roots of Ahypo plants was also in accordance with studies showing that Suc-limiting conditions, led a the induction of the *AtTPS8-AtTPS11* genes in Arabidopsis (Baena-González *et al.*, 2007, Ramon *et al.*, 2009). Also relevant to the above results is the finding that the overexpression of *OsTPS9* in rice significantly increased tolerance toward cold and salinity stress through its proposed association with OsTPS1 (Li *et al.*, 2011; Zang *et al.*, 2011). In contrast, the general WDS-unresponsiveness of *class II TPS*, *TPP* and *TRE* genes in Acau and Ahyb could have contributed to their WDS sensitivity.

Conversely, the differential Pro and RFOs accumulation observed during WDS and R strongly suggests their participation as WDS tolerance factors in amaranth. This proposal is supported by the known role of these compounds as osmoregulators, antioxidants, ROS scavengers, signaling molecules and/ or as C reservoirs for post-stress recovery (Reguera *et al.*, 2013; ElSayed *et al.*, 2014; Kaur *et al.*, 2015; Bascuñán-Godoy *et al.*, 2016).

WDS was also observed to influence the expression levels of RFO biosynthetic genes in a differential way. The differences observed between WDS tolerant and susceptible amaranths were mostly quantitative and were of importance in roots, where the expression *AhGolS1* and *AhRafS* was significantly higher in WDS-tolerant species, especially under SWDS (Table 4). These results were consistent with findings in leaves of *Coffea canephora* clones with contrasting tolerance to WDS, where the expression of the *CcGolS1* gene differed between drought-tolerant and -sensitive clones, being strongly repressed in the latter (dos Santos *et al.*, 2015). Additionally, a related study in *C. arabica* reported that, similar to amaranth, the *CaGolS1* isoform was highly responsive to WDS (dos Santos *et al*., 2011). Likewise, the results in amaranth agreed with several other studies showing that the expression of *GolS* genes was congruous with abiotic stress tolerance in Arabidopsis (Taji *et al.*, 2002; Nishizawa *et al.*, 2008) and in transgenic tobacco plants (Kim *et al.*, 2008; Wang *et al.*, 2009). However, similar to observations in *C. canephora*, higher expression levels of these genes in amaranth leaves and roots did not always coincide with an accumulation of their respective RFOs. Such was the case of Gol, whose amounts were decreased or were undetectable in leaves of Ahypo and in roots of both WDS-susceptible amaranths. Likewise, decreased or unchanged Raf root content in SWDS, and the accumulation of Sta and Ver in leaves of stressed amaranth plants, were contrary to their corresponding gene expression patterns.

Nevertheless, WDS tolerance and RFOs accumulation in amaranth were in agreement with high leaf and root contents of Raf under MWDS and to foliar accumulation MI, Gol and Raf under SWDS, in Ahypo and Acru, respectively. Interestingly, the observed MI buildup may have supplied additional osmoregulatory and antioxidant activity, as previously reported (Ishitani *et al.*, 1996; Duan *et al.*, 2012). Similar results were reported in *Chenopodium quinoa*, an amaranth close relative (Downie *et al.*, 1997). Thus, an increase in MI and/ or Raf levels was observed in leaves of two contrasting *C. quinoa* genotypes subjected to WDS (Bascuñán-Godoy *et al.*, 2016). However, contrary to Ahypo and Acru, Raf levels accumulated in R, which, in quinoa, was proposed to act as a C reservoir utilized for post-stress recovery (Karner *et al.*, 2004). Amaranth RFO accumulation patterns in response to WDS were also similar to those reported in a WDS tolerant alfalfa cultivar able to accumulate Raf and Gol in roots during stress (Kang *et al.*, 2011). A greater accumulation of shoot flavonoids and isoflavonoids was also proposed to contribute to higher WDS in alfalfa. The latter was in accordance with a previous report showing that WDS induced the accumulation of betacyanins and the induction of betacyaninbiosynthetic genes in vegetative tissues of Ahypo cultivars (Casique-Arroyo *et al.*, 2012).

The observed accumulation of Sta and Ver in roots of WDS-stressed amaranth plants was similar to that reported in leaves of *C. arabica* (dos Santos *et al.*, 2011), although it did not seem to affect WDS tolerance in amaranth. On the other hand, other results (Fig. S4-S9), suggest that putative RFOs with a higher degree of polymerization differentially accumulated in leaves of Ahypo and Acru and may have, therefore, contributed to their WDS tolerance. This possibility remains to be determined. Nevertheless, it was in agreement with dos Santos *et al.* (2011, 2015) who argued that the drought-related increase in Gol biosynthesis in coffee was funneled to the generation of larger stress-protective RFOs by unidentified glycosyltransferases.

On the other hand, WDS tolerance in Ahypo and Acru was also defined by significantly higher Pro contents in leaves, principally during SWDS. Significantly higher Pro amounts also accumulated in Ahypo leaves during MWDS (Fig. 4A). On the other hand, Pro accumulation in roots in response to SWDS was, in general, similar in all species (Fig. 4B). Likewise to the behavior observed in alfalfa (Kang *et al.*, 2011), but contrary to the pattern reported in quinoa (Razzaghi *et al.*, 2015; Bascuñán-Godoy *et al.*, 2016), Pro levels declined during R in all species, except in roots of Ahyb. This display coincided with several studies reporting its rapid metabolism in order to provide N and reducing power during stress recovery processes (Hayat *et al.*, 2012; Kaur *et al.*, 2015). Conversely, the significantly higher Pro amounts additionally detected in Ahyb roots during R might partly explain the remarkable recovery observed when severely dehydrated Ahyb plants were re-watered. Pro accumulation was also found to be a contributing factor to WDS tolerance in quinoa (Razzaghi *et al.*, 2015; Bascuñán-Godoy *et al.*, 2016) and alfalfa (Kang *et al.*, 2011).

The characteristic modifications in NSC contents that occur in plants under WDS, both in response to reduced photosynthesis and to the need to maintain water uptake and cell turgor (Seki *et al.*, 2007; Pinheiro and Chaves, 2010) were also observed in amaranth. However, the distinct patterns observed between species suggested that they might have contributed to their different WDS tolerance. Thus, tolerance in Ahypo was associated with inherently low foliar starch levels than became even lower in stressed plants. On the other hand, it presented a basal high Hex/ Suc ratio in roots, which remained practically unchanged by posterior treatments and also underwent a strong depletion of starch levels during WDS and R. The above also suggest that WDS-responsive root CWI, VI, SuSy and amylase enzymes may have been contributing factors to the WDS tolerance observed in Ahypo. Intermediate Acru shared with Ahypo the strong stress-related depletion of starch reserves in both leaves and roots, whereas sensitive Acau and Ahyb had NSC patterns that were essentially the opposite of those observed in Ahypo. Previous reports showing that severe defoliation led to a drastic reduction of C reserves and the induction of various sucroytic and amylolytic enzyme genes (Cisneros, 2016; Castrillón Arbeláez *et al.*, 2012), including two of the four SuSy genes present in the grain amaranth genome (Clouse *et al.*, 2016) advocate this proposal.

The Ahypo NSC fluctuation observed in roots was consistent with the C flow from starch and/ or Suc to Hex triggered as an osmotic adjustment response to WDS in rice and other plants. It agreed, as well, with the increase in invertases and SuSy gene expression and activity that led to an accumulation of Hex in rice plants subjected to WDS (Reguera *et al.*, 2013).

On the other hand, the opposite behavior was observed in WDS susceptible Acau and Ahyb, in which Suc levels tended to increase and starch reserves were less severely depleted during WDS and R. This supports the proposal that varying patterns of NSC accumulation are an additional WDS-tolerance contributing factor in amaranth. However, other aspects not explored in this study, such as fluctuations in N partitioning, have been found to be conductive to WDS tolerance in closely related species, such as quinoa (Bascuñán-Godoy *et al.*, 2016).

Another evident difference detected between WDS-tolerant and WDS-susceptible amaranths, was the variable expression of the ABA marker genes (Tables 5, 6). This suggests that differences in ABA content and/ or sensitivity could be additional factors contributing to the differential WDS tolerance observed in amaranth, as previously described in alfalfa (Kang *et al.*, 2011). In this respect, the general unresponsiveness to WDS of the SnRK2 subgroup of genes analyzed in this study was intriguing (Supplementary Tables S2, S3), considering that *SnRK2* genes are considered to play an important role in stress amelioration, partly through their involvement in the ABA signaling pathway (Liu *et al.*, 2013; Lind *et al.*, 2015)

**Table 6.**
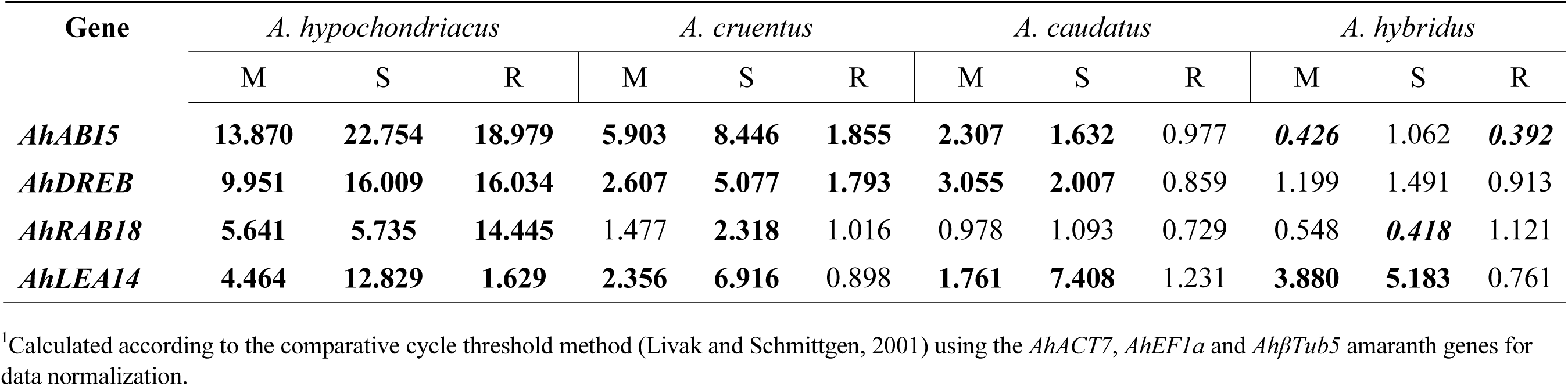
Relative expression values^1^ of abscisic acid (ABA) marker genes in roots of four *Amaranthus* species subjected to two levels of water-deficit stress (moderate [**M**] and severe [**S**]) and to subsequent recovery ([**R**]). Induced (normalized expression values ≥ 2.0; in normal text) and repressed (normalized expression values ≤ 0.5; in italicized text) expression values are emphasized in bold.

In conclusion, this study revealed that differential WDS tolerance between grain amaranth species and leafy, undomesticated, Ahyb, was due to multiple factors. Contributing factors to the improved WDS tolerance observed in Ahypo and Acru, were augmented levels in leaves and/ or roots of Pro and Raf. The WDS-accumulation of Raf in leaves of these species was consistent with augmented *AhGolS1* and *AhRafS* expression levels. Unknown compounds, possibly structurally related to RFOs, were also found to differentially accumulate in leaves of WDS-tolerant species. Additionally, high Raf/ Ver ratios in leaves were found to be a possible determinant of WDS tolerance in amaranth. Moreover, clearly contrasting NSC patterns of accumulation/ depletion in response to WDS and R were observed in leaves and roots of WDS-tolerant and WDS-susceptible amaranth plants. Thus, high Hex/ Suc ratio in roots correlated with superior WDS-tolerance in Ahypo, which was in accordance with the induced activity of CWI, VI and SuSy in response to WDS. A severer depletion of starch reserves, which coincided with significantly increased amylase activity in roots, together with lower soluble NSCs in leaves, also appeared to correlate with WDS-tolerance in amaranth. This, in addition to higher basal levels of Hex in roots of Ahypo, which became even higher in response to WDS. Also significant was the high expression levels of ABA-marker genes in Ahypo plants, which suggested that the WDS tolerance shown by this species could be linked to a higher responsiveness to ABA-related WDS-tolerance mechanisms. Finally, the induced expression of *AhTRE* expression in leaves and of *AhTPS9*, *AhTPS11*, *AhGolS1 and AhRafS* in roots could be employed as markers of WDS tolerance in amaranth.

## Abbreviations

*ABA*: (abscisic acid)
*Ahypo*: (*Amaranthus hypochondriacus*)
*Acru*: (*A. cruentus*)
*Acau*: (*A. caudatus*)
*Ahyb*: (*A. hybridus*)
*CWI*: (cell wall invertase)
*CI*: (cytoplasmic invertase)
*Glu*: (glucose)
*Gol*: (galactinol)
*GolS*: (galactinol synthase)
*Fru*: (fructose)
*MWDS*: (moderate water deficit stress)
*NSC*: (nonstructural carbohydrate)
*Pro*: (proline)
*Raf*: (raffinose)
*RafS*: (raffinose synthase)
*RFO*: (Raffinose Family Oligosaccharides)
*Sta*: (stachyose)
*R*: (recovery)
*Suc*: (sucrose)
*RT*: (retention time)
*SuSy*: (sucrose synthase)
*SWDS*: (severe water-deficit stress)
*TF*: (transcription factor)
*T6P*: (trehalose-6-phosphate)
*TPS*: (trehalose-6-phosphate synthase)
*TPP*: (trehalose phosphate phosphatase)
*Tre*: (trehalose)
*TRE*: (trehalase)
*VI*: (vacuolar invertase)
*WDS*: (water deficit stress)

## Supplementary data

**Fig. S1.** Phylogenetic analysis of amaranth class I and II trehalose phosphate synthase proteins.

**Fig. S2.** Phylogenetic analysis of amaranth trehalose phosphate phosphatase proteins.

**Fig. S3.** Phylogenetic analysis of the amaranth trehalase protein.

**Fig. S4.** Accumulation of unidentified RFO-like compounds during WDS and R in leaves of amaranth plants.

**Fig. S5.** Accumulation of unidentified RFO-like compounds during WDS and R in roots of amaranth plants subjected to WDS.

**Fig. S6.** TLC separation of soluble NSCs and RFOs accumulating in leaf and roots of amaranth plants subjected to WDS.

**Fig. S7.** HPAEC-PAD traces of Ahypo leaf extracts showing the presence of un-identified RFO-like compounds in control and stressed plants.

**Fig. S8.** HPAEC-PAD traces of Acau roots extracts showing the presence of un-identified RFO-like compounds in control and stressed plants.

**Table S1.** List of qPCR primers used in this study.

**Table S2.** Expression patterns of selected *SnRK1* and *SnRK2* genes in leaves of four amaranth species during WDS and R.

**Table S3.** Expression patterns of selected *SnRK1* and *SnRK2* genes in roots of four amaranth species during WDS and R.

## ACKNOWLEDGEMENTS

Support by “La Coordinadora Nacional de las Fundaciones Produce A.C. (COFUPRO; grant DF00000940), and CONACyT, México (grants 339254 and 371475, to ATGR and ICH, respectively) is acknowledged. The authors are also grateful for the support provided by México Tierra de Amaranto A. C., and The Deborah Presser-Velder Foundation.

